# A quasi-digital approach to peptide sequencing using tandem nanopores with endo- and exo-peptidases

**DOI:** 10.1101/018432

**Authors:** G. Sampath

## Abstract

A method of sequencing peptides using tandem cells (*RSC Adv*., 2015, **5**, 167-171; *RSC Adv*., 2015, **5**, 30694-30700) and peptidases is considered. A double tandem cell (two tandem cells in tandem) has three nanopores in series, an amino-acid-specific endopeptidase attached downstream of the first pore, and an exopeptidase attached downstream of the second pore. The endopeptidase cleaves a peptide threaded through the first pore into fragments that are well separated in time. Fragments pass through the second pore and are each cleaved by the exopeptidase into a series of single residues; the latter pass through the third pore and cause separate current blockades that can be counted. This leads to an ordered list of integers corresponding to the number of residues in each fragment. With 20 cells, one per amino acid type, and 20 peptide copies, the resulting 20 lists of integers are used by a simple algorithm to assemble the sequence. This is a quasi-digital process that uses the lengths of subsequences to sequence the peptide, it differs from conventional analog methods which seek to identify monomers in a polymer via differences in blockade levels, residence times, or transverse currents. Several implementation issues are discussed. In particular the problem of fast analyte translocation, widely considered intransigent, may be resolved through the use of a sufficiently long (40-60 nm) third pore. This translates to a required bandwidth of 1-2 MHz, which is within the range of currently available CMOS circuits.

## 1. Introduction

Nanopores have been investigated over several years for their potential use in the analysis of polymers, in particular DNA [1]. There is now a growing interest in other kinds of polymers such as polyethylene glycol [2] and proteins/peptides [3,4]. Here a scheme for peptide sequencing is described that is based on a modified version of a tandem cell with cleaving enzymes that was proposed and studied earlier [5,6]. A tandem cell consists of two electrolytic cells joined in tandem and has the structure [*cis*1, membrane with upstream nanopore (UNP), *trans*1/*cis*2, membrane with downstream nanopore (DNP), *trans*2]. An enzyme attached to the downstream side of UNP successively cleaves the leading monomer in a polymer that has threaded through UNP; the cleaved monomer then translocates to and through DNP where the resulting ionic current blockade level (and/or other discriminators [6]) is used to identify it. With DNA the enzyme is an exonuclease [5], with peptides it is an amino or carboxy peptidase [6]. The approach does not use labels or immobilization.

The present communication describes a method of peptide sequencing using a double tandem cell (two tandem cells connected in tandem), an endopeptidase to break the peptide into fragments, and an exopeptidase to obtain the lengths of the latter. With 20 such cells these length data are used by a simple algorithm to assemble the sequence.

## 2. Peptide sequencing using tandem cells with endo and exopeptidases: a method based on blockade counts

A double tandem cell is an electrolytic cell with four compartments connected in series and bridged by three nanopores. An amino-acid-specific endopeptidase is attached downstream of the first (or upstream) nanopore (UNP) and an exopeptidase is attached downstream of the second (or middle) nanopore (MNP). A peptide drawn into the first pore by a potential difference is cleaved by the endopeptidase after (or before) every occurrence of the amino acid. (Here, ‘before’ and ‘after’ may be interpreted as ‘at the amino end of’ and ‘at the carboxy end of’ or the reverse.) The resulting fragments pass through MNP and are each cleaved by the exopeptidase on the downstream side of MNP into a series of single residues. The latter, well separated in time, pass through the third (or downstream) nanopore (DNP) and cause ionic current blockades that are measured as distinct pulses with width given by a residue’s residence time in DNP. By counting the number of pulses coming from each fragment, an ordered list of fragment lengths (where fragment length = number of residues in a fragment) is obtained. Length lists from 20 such cells and 20 peptide copies, one for each residue type, are then used by a simple algorithm to assemble the sequence. By using a sufficiently long pore (40-60 nm) the bandwidth can be brought down to 1-2 MHz, which is within the range of currently available CMOS circuits [7]. In this quasi-digital approach, subsequence lengths, rather than analog differences in current amplitudes, residence times, or transverse currents, are used to sequence a peptide. Modified amino acids can be handled the same way as natural amino acids.

The report is summarized as follows: In Section 3 the structure and function of a double tandem cell is briefly described and a procedure to obtain fragment length information from a cell outlined. Section 4 describes an algorithm to assemble the peptide sequence using length lists from 20 such cells and traces through the steps of an illustrative example. In Section 5 the minimum cleaving intervals required between successive cleavings by the endopeptidase and the exopeptidase for accurate sequencing are given. Section 6 looks at the detection bandwidth required and its dependence on pore length and fragment and residue translocation times. Section 7 discusses error detection and correction. Section 8 concludes with a brief discussion of implementation issues. An appendix provides supplementary information, including brief notes on a mathematical model of the tandem cell, details of calculations, additional notes on implementation, and a list of related references.

## 3. A double tandem cell for peptide sequencing

Figure 1 shows a schematic of the double tandem cell. It has the structure [*cis*1, membrane with upstream nanopore (UNP), *trans*1/*cis*2, membrane with middle nanopore (MNP), *trans2*/*cis3*, membrane with downstream nanopore (DNP), *trans*3]. A peptide with a poly-X header (X = one of the charged amino acids: Arg, Lys, Glu, Asp; the charge on X depends on the electrolyte pH) is drawn into UNP by the electric field due to V_07_ (= 60-90 mV), most of which (∼98%) drops across the three pores [1]. An endopeptidase specific to amino acid AA attached downstream of UNP cleaves the peptide after all n (≥ 0) points where AA occurs, resulting in n+1 fragments that translocate to and through MNP. On the downstream side of MNP an exopeptidase (amino or carboxy) successively cleaves every residue in a fragment, these single residues translocate through DNP where they cause current blockades, one per residue. The resulting current record for the cell yields an ordered list of integers corresponding to the number of residues in each fragment.

**Fig. 1.**
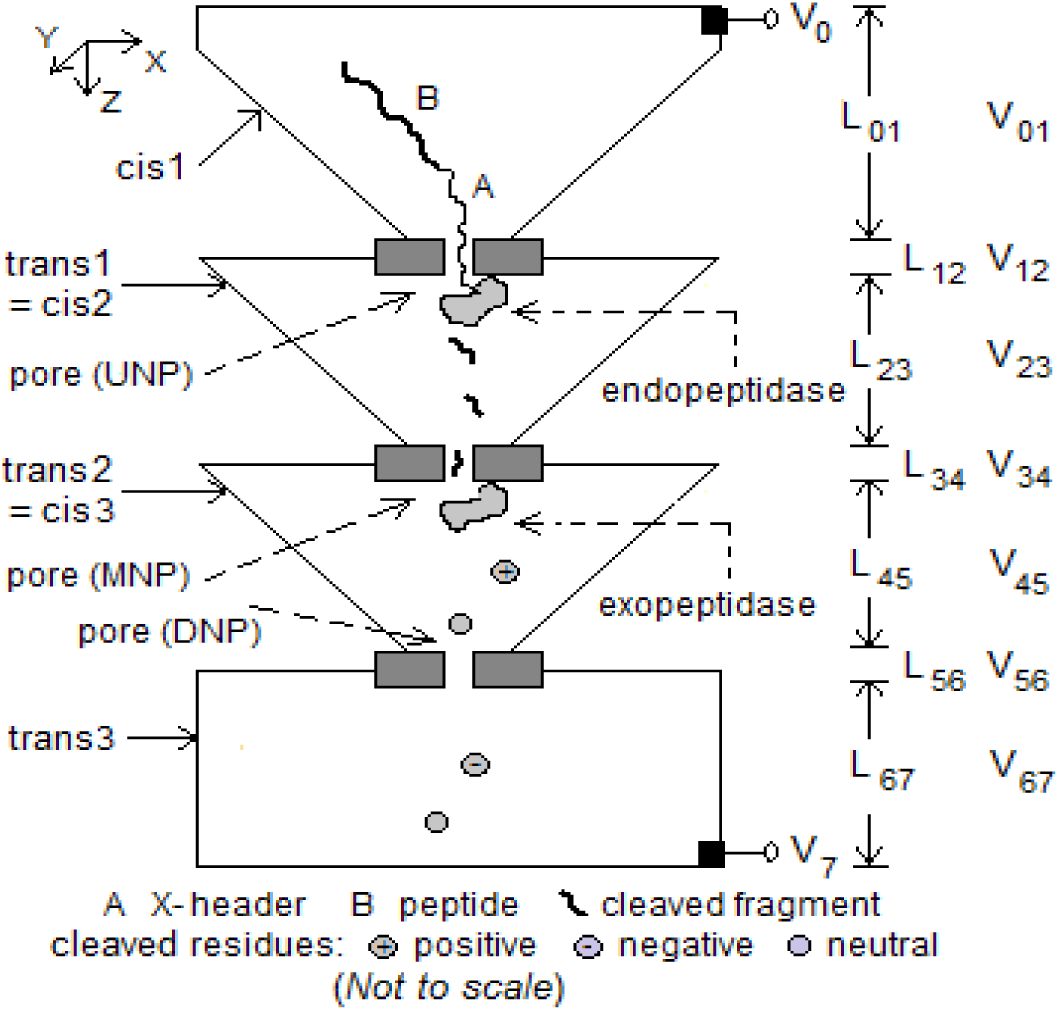
Modified tandem cell for peptide sequencing in 7 stages. Dimensions: 1) *cis*1: box of height 1 μm tapering to 10 nm^2^; 2) membrane with UNP: length 8-10 nm, radius 2-3 nm; 3) *trans*1/*cis*2: box of height 1 μm tapering from 1 μm^2^ to 10 nm^2^; 4) membrane with MNP: length 8-10 nm, radius 2-3 nm; 5) *trans*2/*cis*3: box of height box of height 1 μm tapering from 1 μm^2^ to 20 nm^2^; 6) membrane with DNP: length 40-60 nm, radius 1-2 nm; 7) *trans*3: box of height 1 μm and cross-section 1 μm^2^. Endopeptidase covalently attached to downstream side of UNP; exopeptidase covalently attached to downstream side of MNP. Electrodes at top of *cis*1 and bottom of *trans*3. V_07_ = 60-90 mV.

The basic tandem cell has been modeled mathematically with a Fokker-Planck equation [5]; the results can be applied separately to each compartment or pore in the double tandem cell. More information is provided in the Appendix, where formulas are given for the mean and standard deviation of the translocation time through a section of the cell in terms of the length of the section, the diffusion constant of a charged or uncharged particle, the mobility of a charged particle, and the drift velocity of a charged particle. Other relevant properties such as the hydrodynamic radius of a fragment and the relationship between residue/peptide charge and the pH value of the solution are included. Translocation statistics of peptides are used to estimate the minimum cleaving intervals required of the endopeptidase and the exopeptidase; this is discussed in Section 5 below.

## 4. Sequence assembly from fragment length data

Sequencing with the double tandem cell can be described in the following terms:

1. Each of the 20 cells yields an ordered sequence of integers equal to the numbers of residues in the fragments coming from the corresponding endopeptidase. If a cell generates only one number, the target amino acid is not in the peptide.
2. From these data the peptide is assembled using the following steps:

- replace the fragment lengths from a cell with cumulative lengths (= positions) and the target amino acid identity (= index to a table of the 20 amino acids)
- invert the position-index pairs
- merge the resulting index sequences
- map indexes to amino acids

The following example illustrates the procedure.

*Example*. Let **AA** = [A, R, N, D, C, Q, E, G, H, I, L, K, M, F, P, S, T, W, Y, V] where AA[i] is the i-th amino acid, 1 ≤ i ≤ 20. Consider the peptide KAYTIATRGGATCR. With 8 cells, one for each of the 8 AA types present in the peptide (K, A, Y, T, I, R, G, C), their fragment length lists are: 12:{1, 13}, 1:{2, 4, 5, 3}, 19:{3, 11}, 17:{4, 3, 5, 2}, 10:{5, 9}, 2:{8, 6}, 8:{9, 1,4}, and 5:{13, 1}, where the number before the colon is the index to **AA** of the cell’s target amino acid type. (There are no sequences listed for the 12 AA types that do not occur in the peptide, but see the last sentence of this paragraph.) The last fragment in each list does not contribute to the sequence and is therefore not considered here. Residue position information is computed as 12:{1}, 1:{2, 6, 11}, 19:{3}, 17:{4, 7, 12}, 10:{5}, 2:{8}, 8:{9, 10}, and 5:{13}, where i:{i_1_, i_2_, …} means ‘Amino acid i occurs in positions i_1_, i_2_, … in the peptide’. This list is obtained by replacing a fragment length in the earlier list with the position in the peptide of the last residue in the fragment. The latter is obtained by summing all lengths up to that fragment in the first list. Inverting the target amino acid index and its position in the peptide leads to the sequence of amino acid indices (12, 1, 19, 17, 10, 1, 17, 2, 8, 8, 1, 17, 5), which maps to KAYTIATRGGATC. (The 12 cells for the absent amino acid types will still be used; all generate a single fragment and return a single number equal to the length of the peptide, indicating absence of the target amino acid type.)

Note that the last residue in the peptide sequence (R in the example) cannot be identified, even with the endopeptidase that targets it, because cleaving is assumed to occur after the target amino acid. To resolve this problem, 20 additional cells, each targeting a different one of the 20 amino acid types, can be used to sequence the peptide in reverse and obtain a sequence of fragments in the reverse order. The last residue in the last fragment of a forward sequencing cell will now appear as the first fragment in one of the reverse sequencing cells. A simpler alternative (which would require an extra step) would be to extend the peptide P_x_ with a short poly-Z trailer (with 1 or more Z residues); the last residue B in P_x_ will appear as a fragment in the list from cell B for the extended peptide. The set of fragment lists can be adjusted accordingly.

## 5. Necessary conditions for accurate sequencing

The proposed method relies on counting pulses due to single residues blockading the ionic current through DNP. For sequencing to be accurate the following conditions must be satisfied (see Appendix for the details):

*Condition 1a*: Cleaved residues must arrive at DNP in their natural order. (This is not a strict requirement as residues are being counted, not identified.)
*Condition 1b*: Blockade pulses in DNP must be distinct and therefore well separated in time. Thus there can be no more than one residue in DNP at any time.
*Condition 2a*: Successive fragments translocating from the endopeptidase downstream of UNP must be separated in time so that the pulse count at DNP due to residues from one fragment (cleaved by the exopeptidase after MNP) is not commingled with the count due to residues from the next fragment.
*Condition 2b*: Fragments must arrive at MNP in their natural order. (This is a strict requirement because fragments contribute their lengths to an ordered list.)

From the Appendix, for a maximum peptide length of 20 and a set of typical operating parameters (V_07_ = 90 mV, V_23_ =∼0.3 mV, V_34_ = ∼15 mV, V_45_ = ∼0.3 mV, V_56_ = ∼60 mV, pH = 7.0, L_12_ = 10 nm, L_23_ = 1 μm, L_34_ = 10 nm, L_45_ = 1 μm, L_56_ = 60 nm) the minimum required cleaving intervals for the endopeptidase and the exopeptidase are

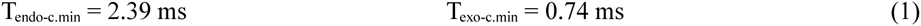

## 6. Translocation time and detection bandwidth

The major factor in the proposed approach is the behavior of cleaved residues (rather than cleaved fragments). The most important consideration is the speed with which a residue translocates through DNP. Translocation times are ordinarily are 10s to 100s of nanoseconds, requiring detection bandwidths that are several tens of MHz. The usually prescribed solution to this is slowing down the residue. There are a variety of methods to do this, most of them have only a marginal effect [8]. Additionally some methods, such as magnetic or optical control, are unusable with peptide fragments or residues, as the latter are intermediate products of internal chemical reactions and cannot be controlled using external agents.

Translocation time (of a fragment or residue) is a function of several parameters: the potential across MNP or DNP, pore (MNP or DNP) length, and the fragment or residue’s diffusion constant D and mobility μ. D is a function of the hydrodynamic radius, which in turn depends on the residues making up the fragment. Mobility is a function of the electric charge on the residue or fragment, which in turn is a function of solution pH. Equations A-1 through A-7 in the Appendix show the relevant relationships. To illustrate the extent of variability, a peptide fragment 20 residues long can have an electric charge anywhere in the range -18q to 18q (q = electron charge), depending on the pH value of the solution.

In the present case the overriding factor is the contribution to the fragment length count coming from a blockade pulse due to a cleaved residue in DNP. If the pulse is wide (or equivalently, the residence time is high) the time resolution required to detect it is decreased; equivalently the required detection bandwidth is also reduced. Thus blockade pulses are wider with longer pores. With a DNP that is 40-60 nm long the pulse width is ∼2 μs (see Table A-3, Columns 7, 9, and 11), and the required bandwidth is 2-4 MHz, which can be measured with currently available CMOS circuits [7]. (Compare this with the usual preference for shorter pores in nanopore-based sequencing, where the goal is better discrimination between successive monomers in a polymer strand [1]. Such a high level of discrimination is not required here because the residues are well separated in time and are being counted rather than identified.)

Table A-3 in the Appendix gives translocation times of single residues in *trans*1/*cis*2 and DNP for three values of pH. Notice the wide difference in times between the more positive residues (R, K, H) and the more negative ones (E, D). The roles are reversed if the sequencing is done with the applied potential reversed. For a given detection bandwidth if the faster residues are missed in one direction they can be picked up with the potential reversed. Thus the reliability of the counting process improves with two fragment lists, one for each of the two directions of applied potential. The error can be minimized over all fragments by experimentally varying the pH and finding the pH value that yields the best results. This can be done independently for each of the 20 cells.

## 7. Error detection and correction

1. *Symbol errors*. These usually arise from incorrect or spurious measurements. A symbol error can be a deletion (a symbol in the sequence is missing) or an insertion (a symbol is sensed in the signal record where there is in reality none). The most common deletion error arises from homopolymers, which are successive monomers that are identical (like the digram GG in the example in Section 4). Homopolymer errors normally cannot occur in the present scheme because by design successive cleaved residues are sufficiently separated in time (see Equation 1 above) when passing through DNP and give rise to distinct pulses. Insertion errors are more likely; they are usually caused by membrane noise. This is discussed next.
2. *Errors due to noise*. Independent of the available bandwidth, noise of one type or another is always present. In an electrolytic cell used for polymer sequencing the type with the most impact is high frequency membrane noise, which arises because the *cis* and *trans* compartments and the membrane in between form a capacitor, as a result of which ions and counterions positioned on either side of the membrane generate random high frequency fluctuations in the pore current [1]. With high detector bandwidth the resulting current spikes increase the probability of false positives (that is, a noise spike is incorrectly counted as a residue), leading to incorrect fragment lengths in the output. A bandpass filter may be used to eliminate spikes outside the band, the required passband can be determined from the mean translocation time of the fastest cleaved residue passing through DNP (see Table A-3 in the Appendix).

Error detection and correction may be based on:

a. standard methods [9], which can be combined to greater effect with the abundant redundancy found in the 20 or more fragment lists;
b. segmentation methods that use hidden Markov models to identify transitions between noise and signal [10];
c. database identification methods that are commonly used in mass spectrometry [11]; they make use of algorithms like Peptide Sequence Tags and Sequest and can be adapted to the present purpose.

## 8. Some implementation considerations

There are a number of factors to consider in implementation, and several parameters may be used to arrive at an optimum design. They include choice of pores, applied voltage, translocation speed, solution pH, detector bandwidth, membrane noise, peptide length, choice of electrolyte, and temperature. Some of these issues are considered next.

1. *Choice of pores*. As discussed in Section 6, a long DNP is necessitated by the need to decrease the detection bandwidth. Synthetic pores made of silicon compounds, such as silicon nitride (Si_3_N_4_), are well-suited for this purpose. Typically they have a length of 30-100 nm, but the fabrication process makes them double-conical rather than cylindrical [12], which would require design parameters to be adjusted suitably. The purpose of UNP and MNP is mainly to feed a peptide or peptide fragment to the associated peptidase for cleaving. Biological pores like alpha hemolysin (AHL) may be better suited because of the need to covalently attach (or biologically fuse) the peptidase to the downstream side of the pore.
2. *Order of fragment entry into MNP*. With a narrow pore, it is possible for a cleaved fragment in *trans*1/*cis*2 translocating single file through a narrow MNP to enter it from the wrong end (C-terminal if the exopeptidase is amino, or N-terminal if carboxy). One way to resolve this problem is to use a wider pore and position the exopeptidase so that it can bind to the fragment and successively cleave the residues from the correct end. The larger width may not have any adverse effects because MNP’s role is only to serve as a conduit for the fragment to be drawn processively to the exopeptidase for the purpose of cleaving.
3. *Current levels and signal-to-noise ratio (SNR)*. The measured quantity of interest is the ionic current through DNP; its value ∼50-60 picoamperes (based on pore conductances of ∼1-2 nS [12,13]). A commonly used method of increasing SNR is to use a bandpass filter. Higher voltages, even if they increase the SNR, can cause problems in sequence order; see Section 3 in the Appendix. For further improvement, error correction methods may be used. (Also see the last paragraph of Section 6.)
4. *Entropy barriers*. It is assumed that the entropy barrier [1] faced by a fragment during its entry into MNP is negligible. When not negligible, it can still be taken into account. Thus the minimum cleaving interval in Equation 1 may be increased to allow for the additional time taken by a fragment to overcome the barrier. The tapers shown in Figure 1 also help lower the entropy barrier for entry of the peptide into UNP, fragment into MNP, and residue into DNP.
5. *Solution pH*. Solution pH plays an important role for two reasons: a) the charge carried by a fragment, which is highly variable and not known in advance, is a function of pH; this is unlike with DNA, where all nucleotide types have approximately the same charge of -q; b) its effect on enzyme reaction rates. The choice of pH is therefore a tradeoff between enzyme efficiency and the need to control translocation speeds. The optimum pH value is best determined by experiment.
6. *Modified amino acids*. The proposed device can be used to sequence peptides containing modified amino acids; this only requires the use of an additional cell with an endopeptidase and peptide copy for each modified amino acid type. (Information on peptidases is available in the MEROPS database, which is accessible at <u>http://merops.sanger.ac.uk</u>. Also see recent review [14].)

For other issues, see ‘Additional Notes’ in the Appendix.

## Appendix

### A-1 Translocation statistics of tandem cell

Following [5], the mean E(T) and variance σ^2^(T) of the translocation time T over a channel of length L that is reflective at the top and absorptive at the bottom with applied potential difference of V are given by

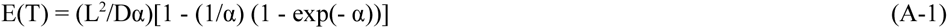

and

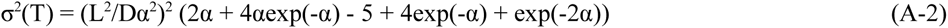

where

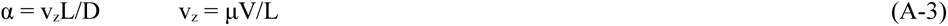

Here v_z_ is the drift velocity due to the electrophoretic force experienced by a charged particle in the z direction, which can be 0, negative, or positive. For v_z_ = 0, these two statistics are

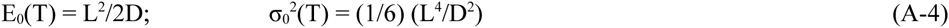

If each section in the double tandem cell is considered independently these formulas can be applied to all the relevant sections: *trans*1/*cis*2 (T = T_*trans*1/*cis*2_; L = L_23_), MNP (T = T_MNP_; L = L_34_), *trans2*/*cis3* (T = T_*trans*2/*cis*3_; L = L_45_), DNP (T = T_DNP_; L = L_56_), and *trans*3 (T = T_*trans*3_; L = L_67_). For an analysis of behavior at the interface between two sections see [5,6].

### A-2 Dependence of peptide translocation on solution pH, charge, diffusion constant, and mobility

Equations A-1 through A-4 involve a number of physical-chemical properties of amino acids: electrical charge (itself dependent on solution pH [15]), hydrodynamic radius, diffusion constant, and mobility. The following paragraphs provide a quantitative description of this dependence and allow calculation of fragment properties as they apply to peptide sequencing in a tandem cell with endopeptidase and exopeptidase. In particular this information is used in the next section to derive a required condition for effective sequencing.

1. The electrical charge carried by a peptide (fragment) P_x_ can be calculated with the Henderson-Hasselbach equation. Let the set of amino acids be **AA** = [A, R, N, D, C, Q, E, G, H, I, L, K, M, F, P, S, T, W, Y, V] where AA[i] is the i-th amino acid, 1 ≤ i ≤ 20. Let the pH value of the solution (electrolyte) be p, kC = kA value of the carboxy end = 9.69, kN = kA value of the amino end = 2.34, N_X_ the number of times residue X (= R, H, K) occurs in the peptide, N_Z_ the number of times residue Z (= D, C, E, Y) occurs, and kX and kZ the kA values of X and Z respectively. kA values are given by the following table:

**Table A-1.**
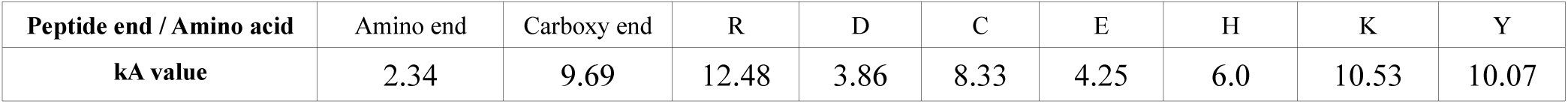 The charge multiplier C_Px_ for peptide P_x_ is given by

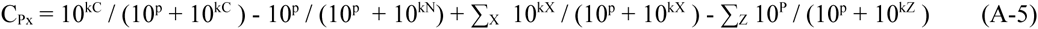

where the summations are over the N_X_ and N_Z_ occurrences of X and Z respectively in P_x_.
2. The hydrodynamic radius R_Px_ of peptide P_x_ = X_1_ X_2_ … X_N_ is obtained recursively as follows:

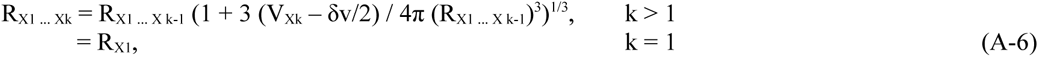

where V_Xk_ and and δv are the van der Waals volumes of X_k_ and a single molecule of water. Hydrodynamic radii of individual amino acids are given in [16] and van der Waals volumes in [17] (both sets of values are reproduced in the Supplement to [6]). This formula holds for small peptides (up to ∼20 residues).
3. The diffusion constant and mobility of P_x_ are given by

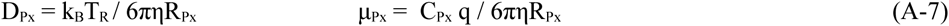

Here k_B_ is the Boltzmann constant (1.3806 × 10^-23^ J/K), T_R_ is the room temperature (298 ° K), η is the solvent viscosity (0.001 Pa.s), and q is the electron charge (1.619 × 10^−19^ coulomb).

As an example, consider peptide P_x_ = RRRRRRRRRRRRRRRRRRKE in the cell with target amino acid K. It translocates through UNP to *trans*1/*cis*2 where it is cleaved by a K-specific endopeptidase into two fragments, F_1_ = RRRRRRRRRRRRRRRRRRK and F_2_ = E. Translocation statistics for the two fragments are calculated at three values of pH (3.0, 7.0 = physiological pH, and 11.0) using Equations A-1 through A-7 for a *trans*1/*cis*2 of height 1 μm and MNP of length 8 nm and radius 2 nm and given below in Table A-2. Because of the high charge multiplier, the applied potential V_07_ is set to ∼80 mV. This results in ∼15 mV across MNP, which is lower than usual in nanopore sequencing (in most studies it is typically ∼100-200 mV). This is because otherwise translocation times for positively charged fragments, especially long ones like F_1_ above, can be excessive. (The P_x_ chosen above is a worst-case example that is used to determine the minimum interval required between two successive cleavings by the endopeptidase; see Section A-3.)

**Table A-2.**

| Peptide fragment | Translocation time in *trans1/cis2* ($10^{-3}$ s) | | | | | | Translocation time in MNP ($10^{-6}$ s) | | | | | |
| --- | --- | --- | --- | --- | --- | --- | --- | --- | --- | --- | --- | --- |
|  | pH = 3.0 |  | pH = 7.0 |  | pH = 11.0 |  | pH = 3.0 |  | pH = 7.0 |  | pH = 11.0 |  |
|  | Mean | Std dev | Mean | Std dev | Mean | Std dev | Mean | Std dev | Mean | Std dev | Mean | Std dev |
| F<sub>1</sub> = RRRRRRRRRRRRRRRRRRK | 0.5324 | 0.4412 | 0.5321 | 0.4408 | 0.5272 | 0.4360 | 147.5404 | 147.5239 | 135.2212 | 135.2046 | 46.1967 | 46.1787 |
| F<sub>2</sub> = E | 0.1799 | 0.1470 | 0.1791 | 0.1461 | 0.1785 | 0.1455 | 0.0472 | 0.0387 | 0.0382 | 0.0300 | 0.0326 | 0.0245 |

*trans1/cis2*: height = 0.5 μm radius = 0.5 μm $V_{23}$ = 0.3mV
MNP: height = 8nm radius = 2 nm $V_{34}$ = 15 mV

Similar calculations for translocation times of single residues through *trans*2/*cis*3 and DNP lead to the results in Table A-3.

**Table A-3.**

| Amino acid | Translocation time in *trans2/cis3* ($10^{-3}$ s) | | | | | | Translocation time in DNP ($10^{-6}$ s) | | | | | |
| --- | --- | --- | --- | --- | --- | --- | --- | --- | --- | --- | --- | --- |
|  | pH=3 |  | pH=7 |  | pH=11 |  | pH=3 |  | pH=7 |  | pH=11 |  |
|  | Mean | Std dev | Mean | Std dev | Mean | Std dev | Mean | Std dev | Mean | Std dev | Mean | Std dev |
| A | 0.1526 | 0.1246 | 0.1523 | 0.1244 | 0.1512 | 0.1233 | 2.5355 | 2.1271 | 2.1903 | 1.7878 | 1.1808 | 0.8193 |
| R | 0.2081 | 0.1702 | 0.2078 | 0.1699 | 0.2062 | 0.1683 | 9.3647 | 8.7947 | 7.6052 | 7.0346 | 3.0031 | 2.4576 |
| N | 0.1709 | 0.1396 | 0.1707 | 0.1394 | 0.1694 | 0.1381 | 2.8405 | 2.3830 | 2.4538 | 2.0029 | 1.3229 | 0.9178 |
| D | 0.1730 | 0.1413 | 0.1716 | 0.1399 | 0.1704 | 0.1387 | 2.6075 | 2.1482 | 1.3069 | 0.8990 | 0.8548 | 0.4994 |
| C | 0.1640 | 0.1340 | 0.1637 | 0.1337 | 0.1613 | 0.1313 | 2.7261 | 2.2871 | 2.2752 | 1.8441 | 0.8102 | 0.4735 |
| Q | 0.1852 | 0.1513 | 0.1850 | 0.1510 | 0.1836 | 0.1497 | 3.0788 | 2.5829 | 2.6597 | 2.1709 | 1.4339 | 0.9948 |
| E | 0.1800 | 0.1470 | 0.1784 | 0.1455 | 0.1771 | 0.1442 | 2.8642 | 2.3841 | 1.3595 | 0.9354 | 0.8888 | 0.5192 |
| G | 0.1331 | 0.1087 | 0.1329 | 0.1085 | 0.1319 | 0.1075 | 2.2114 | 1.8552 | 1.9103 | 1.5593 | 1.0299 | 0.7146 |
| H | 0.2017 | 0.1650 | 0.2000 | 0.1633 | 0.1984 | 0.1618 | 9.0679 | 8.5153 | 3.0883 | 2.5562 | 1.5493 | 1.0749 |
| I | 0.1858 | 0.1518 | 0.1856 | 0.1515 | 0.1842 | 0.1502 | 3.0883 | 2.5909 | 2.6679 | 2.1776 | 1.4383 | 0.9979 |
| L | 0.1944 | 0.1588 | 0.1942 | 0.1585 | 0.1927 | 0.1571 | 3.2313 | 2.7109 | 2.7914 | 2.2785 | 1.5049 | 1.0441 |
| K | 0.2133 | 0.1745 | 0.2130 | 0.1742 | 0.2102 | 0.1714 | 9.5988 | 9.0145 | 7.7928 | 7.2080 | 1.8892 | 1.3710 |
| M | 0.1766 | 0.1443 | 0.1764 | 0.1440 | 0.1751 | 0.1428 | 2.9358 | 2.4630 | 2.5361 | 2.0701 | 1.3673 | 0.9486 |
| F | 0.1921 | 0.1569 | 0.1919 | 0.1567 | 0.1904 | 0.1553 | 3.1932 | 2.6789 | 2.7585 | 2.2516 | 1.4871 | 1.0318 |
| P | 0.1537 | 0.1255 | 0.1535 | 0.1253 | 0.1524 | 0.1242 | 2.5545 | 2.1431 | 2.2068 | 1.8013 | 1.1897 | 0.8254 |
| S | 0.1583 | 0.1293 | 0.1581 | 0.1291 | 0.1569 | 0.1279 | 2.6308 | 2.2071 | 2.2726 | 1.8550 | 1.2252 | 0.8501 |
| T | 0.1744 | 0.1424 | 0.1741 | 0.1422 | 0.1728 | 0.1409 | 2.8977 | 2.4310 | 2.5032 | 2.0432 | 1.3495 | 0.9363 |
| W | 0.2007 | 0.1639 | 0.2005 | 0.1637 | 0.1990 | 0.1622 | 3.3361 | 2.7989 | 2.8820 | 2.3524 | 1.5537 | 1.0780 |
| Y | 0.2047 | 0.1672 | 0.2045 | 0.1669 | 0.2015 | 0.1641 | 3.4029 | 2.8548 | 2.9377 | 2.3975 | 1.0521 | 0.6253 |
| V | 0.1904 | 0.1555 | 0.1901 | 0.1553 | 0.1887 | 0.1539 | 3.1646 | 2.6549 | 2.7338 | 2.2314 | 1.4738 | 1.0226 |

*trans2/cis3*: height = 0.5 μm radius = 0.5 μm $V_{23}$ = 0.6mV
DNP: height = 60 nm radius = 2 nm $V_{34}$ = 60 mV

### A-3 Conditions for ordered fragment entry into MNP and single residue occupancy in DNP

The proposed method relies on counting pulses due to single residues blockading the ionic current through DNP. Therefore for sequencing to be accurate the following conditions must be satisfied.

*Condition 1a*: Blockade pulses in DNP must be distinct and therefore well separated in time. Thus there can be no more than one residue in DNP at any time.

*Condition 1b*: Cleaved residues must arrive at DNP in their natural order.

*Condition 2a*: Fragments must arrive at MNP well separated in time so that the pulse count at DNP for the residues of one fragment is distinct from the pulse count for the residues of the next fragment.

*Condition 2b*: Fragments must arrive at MNP in their natural order. (This is a strict requirement because fragments contribute integers to an ordered list.)

Each of these conditions sets a minimum time for the interval between successive cleavings by the two enzymes. Let T_exo-c-min_ and T_endo-c-min_ be the minimum cleaving intervals of the two enzymes that satisfy the above four conditions. Assume that the translocation times have distributions with 6σ support (σ is the standard deviation, see Equations A-2 and A-4). Following the procedure in [6] (see Supplement therein), if X_1_ and X_2_ are two residues cleaved in succession by the exopeptidase, *Conditions 1a and 1b* are both satisfied if

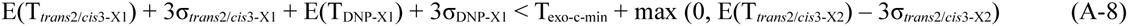

Let pH = 7.0 (physiological pH). From columns 3 and 4 in Table A-3, the second term on the right side of the inequality is 0 for any X2 so that

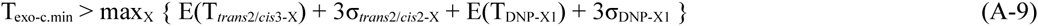

where X ranges over all the 20 amino acids. The maximum translocation times occur for X = K (Lys) both in *trans*2/*cis*3 and in DNP. Using the data for Lys in columns 3, 4, 9, and 10 in Table A-3

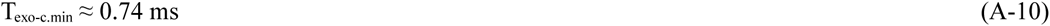

With *trans*1/*cis*2 and MNP the analysis is not as simple; a fragment can have any length and carry a total electric charge that can be positive, negative, or 0. If the peptide length is limited to 20 (this is the optimum length in a high-quality mass spectrometer [11]), a worst case analysis can be done with a maximum fragment length of 19. (When the fragment length is 20, no cleaving has occurred after UNP. The whole peptide passes into MNP and is cleaved into 20 single residues by the exopeptidase; no information about the sequence other than the length is generated in DNP.) The sample peptide P_x_ = RRRRRRRRRRRRRRRRRRKE used in the example calculations above is a worst-case example of a peptide that is broken into two fragments F_1_ = RRRRRRRRRRRRRRRRRRK and F_2_ = E in the cell for K. It can be used to calculate the minimum interval required between successive fragments created by and/or exiting the peptidase for target amino acid X = K.

F_1_ has maximum positive charge and F_2_ has a negative charge. Thus the translocation times in *trans*1/*cis*2 and MNP are maximum for F_1_ and minimum for F_2_. Conditions 2a and 2b are both satisfied if

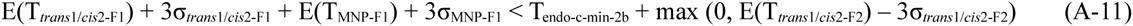

From columns 3 and 4 in Table A-2 the second term in the inequality on the right is 0. Using data from columns 3, 4, 9, and 10,

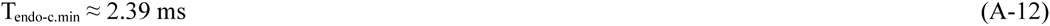

Notice that the solution pH plays an indirect role in the above calculation, it is also a major contributing factor to peptidase functioning. The optimal value of pH to use can be determined by experiment independently for each of the 20 cells.

### A-4 Additional notes

1. *Identifying the last residue in a peptide*. An alternative to the use of reverse sequencing cells to identify the last residue in a peptide is to sequence a copy of the peptide in the forward direction using a second cell with an endopeptidase that cleaves before rather than after the target amino acid.
2. *Order of fragment entry into MNP*. A fragment can enter MNP amino-end first or carboxy-end first. However the order is not important as the information sought is the number of residues, not their identity or sequence.
3. *Order of entry of peptide into UNP*. The assembly algorithm described in Section 4 implicitly assumes that entry of a peptide into UNP in each of the cells is for all of them N-terminal first or all of them C-terminal first. This is a reasonable assumption because of the charged X-header. However, there is a non-zero probability that the peptide may enter wrong end first, so some of the fragment length lists obtained will be in the reverse order. The assembly algorithm needs to be modified to take this into account. Preliminary analysis suggests that the required modification may not be a major one.
4. *Independence of cells*. Each cell targets a different amino acid and operates independent of the other cells. This means that the cell can be independently optimized for enzyme reaction rates, applied voltage, pH value, etc.
5. *Effect of the X-header*. A charged X-header of length M attached to the target peptide is used to induce the peptide to enter UNP. As it is itself cleaved by the endopeptidase for X it generates M additional fragments containing a single X residue; the output of the X cell has to be adjusted accordingly. Similarly the length of the first fragment going into DNP also needs to be adjusted.
6. *Entropy barrier at entrance to MNP.* As noted earlier, a fragment entering MNP may face an entropy barrier [1]. For short peptides with lengths up to ∼20, this may not be significant. A longer fragment, however, may be coiled on itself and take more time to enter MNP; this means an increase in the minimum cleaving interval at MNP (see Section A-3). The value in Equation 1 in the main text can be multiplied by a suitable factor to account for barrier effects.
7. *Sticky fragments/residues*. The problem of fragments or residues sticking to pore or compartment walls may be resolved through the use of non-stick additives [18] or wall coatings [19].
8. *Whole protein sequencing*. If fragment lengths created by the cleaving enzyme are not large <20 a folded protein could be loaded into the tandem cell and unfolded by an unfoldase enzyme like ClpX [20] before cleaving and sequencing.
9. *Hafnium oxide pores*. Recent studies using high bandwidth (∼1 MHz) detectors have shown that a HfO_2_ membrane < 10 nm long can slow down translocating DNA molecules [21]. (The slowdown is believed to be due to interactions of the DNA with the walls of the pore.) However, fabrication of such pores appears to require an inordinate amount of time.
10. *Other measurements*. While the proposed method is centered on extracting sequence identity from integers representing the lengths of cleaved fragments, other parts of the current record (blockade levels, blockade pulse widths, translocation times, etc.) can be used for such purposes as obtaining higher-order correlations, correcting sequence assembly errors, etc. They may also be used in bioinformatics applications.
11. *Applicability of the proposed approach to DNA sequencing*. At present the question whether this approach can be applied to DNA sequencing has to be answered in the negative because nucleotide-specific endonucleases, one for each of A, T, C, and G, are required. Known endonucleases cleave a strand at a scission point without distinguishing among base types. If nucleotide-specific endonucleases can be found or synthesized then the approach presented here may simplify DNA sequencing considerably.

For other implementation-related issues affecting tandem cells see discussions in [5,6]. For a review of nanopore fabrication methods see [22].

